# Ultrahigh throughput evolution of tryptophan synthase in droplets via an aptamer-biosensor

**DOI:** 10.1101/2023.10.11.561886

**Authors:** Remkes A. Scheele, Yanik Weber, Friederike E. H. Nintzel, Michael Herger, Tomasz S. Kaminski, Florian Hollfelder

## Abstract

Tryptophan synthase catalyzes the synthesis of a wide array of non-canonical amino acids and is an attractive target for directed evolution. Droplet microfluidics offers an ultrahigh throughput approach to directed evolution (>10^7^ experiments per day), enabling the search for biocatalysts in wider regions of sequence space with reagent consumption minimized to the picoliter volume (per library member). While the majority of screening campaigns in this format on record relied on an optically active reaction product, a new assay is needed for tryptophan synthase. Tryptophan is not fluorogenic in the visible light spectrum and thus falls outside the scope of conventional droplet microfluidic read-outs which are incompatible with UV light detection at high throughput. Here, we engineer a tryptophan DNA aptamer into a biosensor to quantitatively report on tryptophan production in droplets. The utility of the biosensor was validated by identifying 5-fold improved tryptophan synthases from ∼100,000 protein variants. More generally this work establishes the use of DNA-aptamer sensors with a fluorogenic read-out in widening the scope of droplet microfluidic evolution.

## Introduction

Directed evolution has enabled the search for proteins that are of immediate interest to the pharmaceutical and chemical industries.^1^ The success of directed evolution is determined by i) the starting point in sequence space, ii) the throughput and library design to probe the surrounding fitness landscape, and iii) the assay through which fitness is determined.^2^ Microtiter plate screening (∼10^3^ variants per day) is readily coupled to a direct measurement of product – either through liquid/gas chromatography and/or mass spectrometry. Cellular or droplet-based ultrahigh throughput screening (uHTS, ∼10^7^ droplets per day) benefits from an increase in throughput but requires an optical signal of the reaction product.^3^

Directed evolution in single cells or droplets has seen the most success using fluorogenic proxy molecules: these small molecules are synthesized with a scissile bond connected to a fluorophore. Upon enzymatic cleavage, the leaving group becomes fluorescent and enables sorting by fluorescence-activated droplet sorting (FADS) sensitively (e.g. >3000 molecules fluorescein per droplet) and rapidly (>kHz).^4^ These substrates are, however, hard to synthesize, and often do not resemble substrates relevant for industrial biocatalysis, e.g. respective chemical challenges are not reflected in activated model substrates. Alternatively, coupled reactions can be used to detect the redox state of cofactors through absorbance-activated droplet sorting (AADS), facilitating the screening of NAD(P)H-dependent transformations.^5–7^ Nevertheless, the bulk of analytes are currently not amenable to any optical assay and remain beyond the reach of uHTS, although mass-activated, dielectrophoretic, and NMR sorting are emerging as a future alternative ^8–10^ (albeit at the price of lower throughput). Simple, fast and modular screening approaches in droplets addressing non-surrogate substrates are highly sought-after and would allow the implementation of a wider spectrum of target reactions in uHTS format for enzyme evolution pipelines.

Engineering protein or oligonucleotide biosensors to detect small molecules of interest, i.e. the reaction product, is an alternative strategy to obviate the need for fluorogenic substrates. Protein biosensors require engineering a conditional inactive state that can be reversed by binding a molecule of interest.^11^ For example, a protein sensor was engineered to detect the redox state of NAD, thus enabling screening of dehydrogenases on non-fluorogenic substrates in droplets by fluorescent activated cell sorting (FACS).^12^

Short oligonucleotides, or aptamers, can be evolved through SELEX to bind any small molecule of interest.^13–15^. Numerous methods exist to engineer the resulting binder into a biosensor suitable for uHTS: RNA-based aptamers can be engineered into a biosensor through fusion with the ‘Spinach’ probe,^14^ which adapts its secondary structure when the original aptamer binds the target analyte, rendering it fluorescent.^16,17^ For DNA-based aptamers, a biosensor can be engineered by fluorescently labelling the aptamer while designing a complementary strand with a quencher.^18^ When the target analyte is present, the complementary strand is displaced in favour of the target analyte, resulting in a fluorescence increase. Recently, the Heemstra group was able to detect unlabelled L-tyrosinamide in droplets using this DNA-based biosensor design for enantiopurity analysis,^19^ setting the scene for future development of label-free assays for evaluating and sorting enzyme variants in droplets.

As such, DNA-aptamers are convenient to use and require only synthesis of the evolved aptamer with a fluorophore and a complementary strand with a quencher to create a biosensor. To utilize DNA aptamers in uHTS, one must (i) find/evolve an aptamer for a small molecule of interest; (ii) ensure that the aptamer does not bind the substrate, i.e. is specific for the product; (iii) manipulate the equilibrium of complementary strand and product binding to create a sensor; (iv) encapsulate the biosensor in single or double emulsion droplets with substrates and unique enzyme variants, and sort by FADS or FACS respectively.

Here, we develop a biosensor from an existing DNA aptamer for L-tryptophan (Trp) **(Figure 1)**. We demonstrate the integration of the Trp biosensor for the uHT evolution of the tryptophan synthase β-subunit from *Pyrococcus furiosus* (from here on referred to as TrpB), which condenses L-serine (Ser) and indole to produce Trp. TrpB has a relaxed substrate scope and has been evolved to produce a wide range of non-canonical amino acids (NCAAs) of industrial interest with good yield.^20^ However, Trp (and its derivatives) cannot be assayed by conventional fluorescence or absorbance-activated droplet sorting,^5^ since any intrinsic optical signal change is too small to be measured at high throughput in microfluidic devices. Assaying ∼100,000 TrpB variants in a single day enabled us to isolate TrpB^A9^, which accumulated four mutations in a single round to make it 5-fold more catalytically efficient than the wildtype TrpB, and establishes the utility of DNA aptamer biosensors for uHTS.

**Figure 1.**
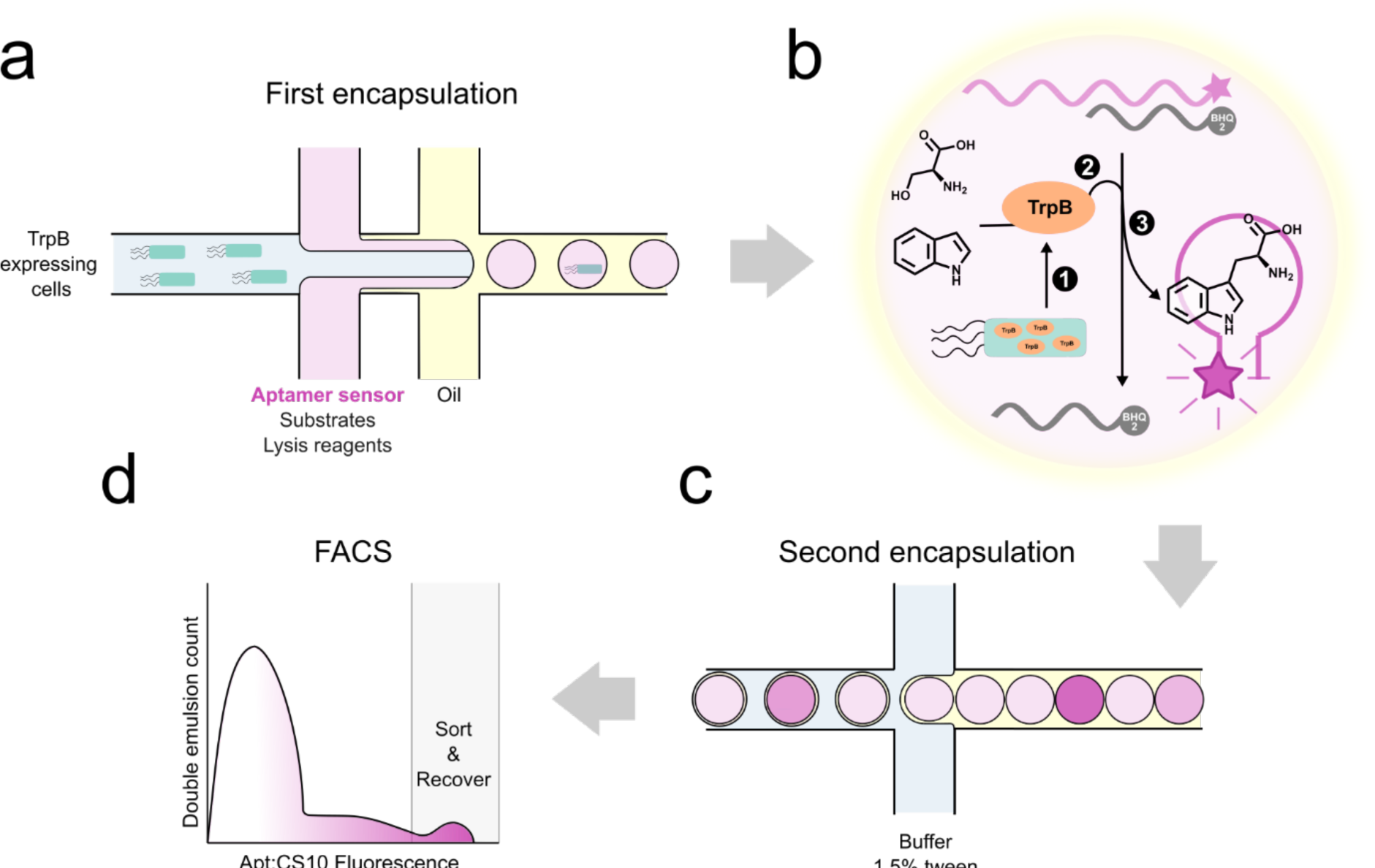
Overview of the screening approach. (a) TrpB-expressing cells are encapsulated in single emulsion droplets with the aptamer sensor, the substrates (Ser, indole) and lysis reagents. (b) 1 - The lysis reagents will release TrpB in each droplet where a cell is present. 2 - TrpB catalysis of Ser and indole to produce Trp. 3 - The fluorescent aptamer is initially self-quenched, but once Trp is bound in favour of the quenching complementary strand, the sensor lights up and becomes fluorescent. The Trp concentration measured as a fluorescence signal is a function of the catalytic efficiency, stability and expression strength of TrpB variants, and screening and selection can be carried out accordingly. (c) The droplets are encapsulated again into double emulsion droplets so that they are compatible with fluorescence-activated cell sorting on a commercial flow cytometer. (d) The genotype from the pool of highly fluorescent droplets is recovered, after which the enriched pool of active variants is rescreened in microtiter plate-based format for single variants of interest. The chip design is shown in **Supplementary Figure 1**.

## Results

### From aptamer to sensor: concentration-dependent fluorescence change upon Trp binding

The starting point for this work was an existing Trp aptamer previously evolved through SELEX with a *K*_d_ of 1.8 µM for Trp.^21^ For the Trp aptamer to work in conjunction with droplet-based screening, a Trp-concentration-dependence of its fluorescent signal is necessary to quantitatively monitor product emergence, in other words, a biosensor for Trp. To create the biosensor, a fluorescently labelled aptamer can be coupled to a complementary strand (CS) with a quencher, which is released in favour of the analyte (**Figure 2A**).^18^ In this case, the secondary structure prediction of the Trp aptamer (**Figure 2A**) suggests the presence of a stem loop (estimated to exist with >99% likelihood).^22^ When the Trp aptamer is bound to a CS, the stem loop would be unable to hybridize, disrupting secondary structure formation, which was thought to disable the aptamers ability to bind Trp. An ideal complementary strand with a quencher (CS) would bind the fluorescent aptamer at saturation when no target analyte is present, so that background fluorescence is minimized. However, the competition between binding of the CS or Trp needs to favour binding of Trp, so that the CS gets displaced with increasing concentrations of Trp and a fluorescent signal emerges.

**Figure 2.**
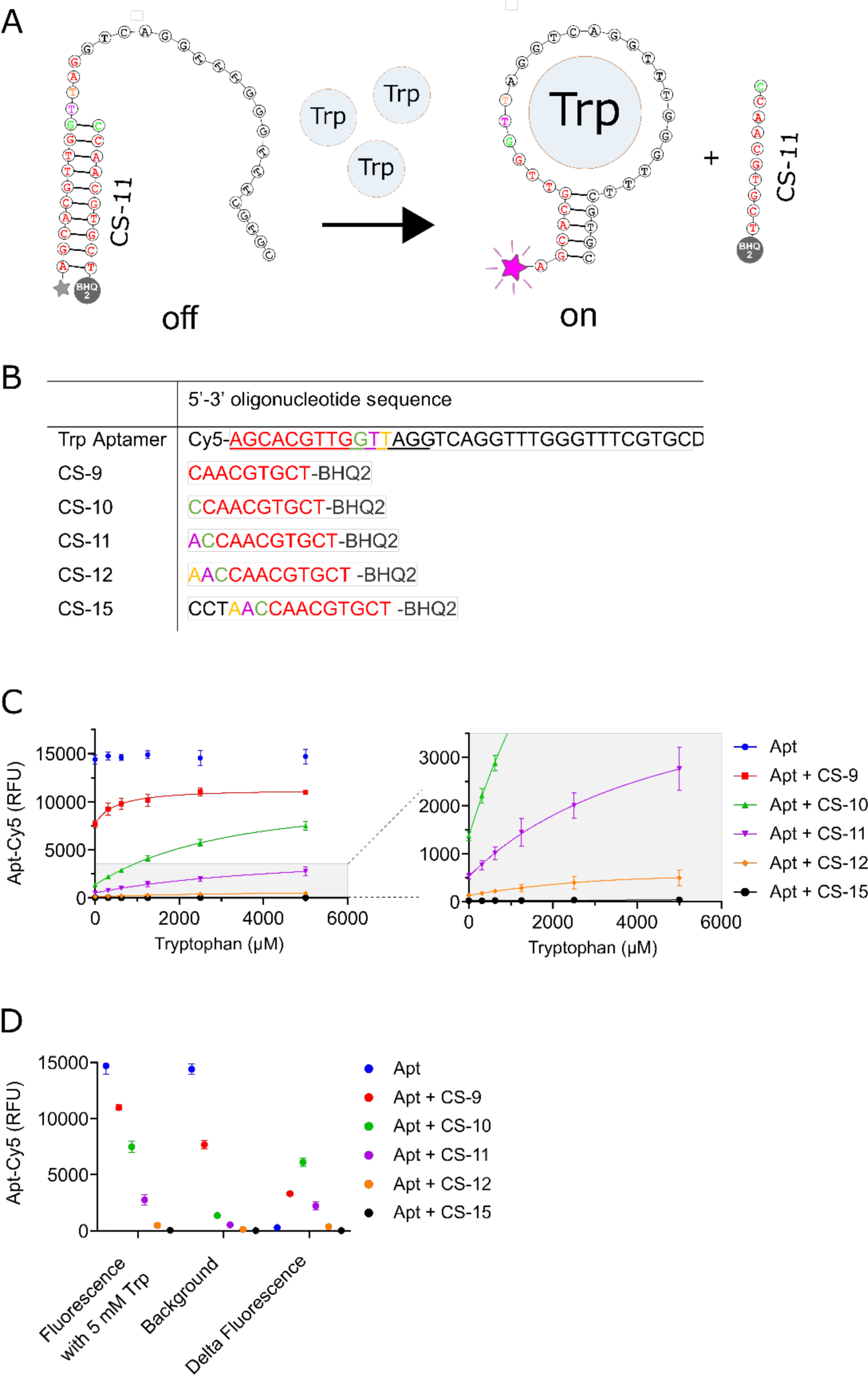
From aptamer to aptamer sensor. (A) The predicted loop structure of the aptamer without CS present (right)^22^ contains a hybridized stem region. Binding of a CS with a quencher (left) puts the BHQ2 in the proximity of the Cy5 fluorescent group on the aptamer, quenching its fluorescence, and prohibits formation of the stem loop. When enough Trp is present to tilt the equilibrium, the CS gets displaced in favour of Trp and the Cy5 fluorophore gets unquenched. (B) The Trp aptamer and CS sequences with Cy5 and BHQ2 moieties respectively, added during synthesis. (C) Titration of Trp to (hybridized) solutions of Trp aptamer without any CS (blue circles) and in the presence of CS-9 (red squares), CS-10 (green triangles), CS-11 (purple triangles), CS-12, (orange diamonds) and CS-15 (black circles). The grey area is enlarged in the second panel (on the right) to show the CS-11, CS-12, and CS-15 titration curves. Standard deviations derived from two technical repeats are shown. (D) The aptamer:CS10 duplex has the highest delta fluorescence in the presence of Trp. Experiments were performed in tryptophan aptamer buffer (TAB), pH 7.4, at 25 °C.

The equilibrium between the aptamer’s affinity for Trp and the aptamer’s affinity for the CS is most easily optimized by altering the length of CS.^18^ Whereas the affinity for the target analyte is a direct outcome of SELEX, the affinity of the CS can be systematically modified based on the dependence of K_D_ on its complementary length to the aptamer (the longer the complementary region, the more stable the duplex) and the entropic assistance of intramolecularity. A panel of CSs were synthesized, altering the length of the CS between 9 and 15 nucleotides (**Figure 2B**). Each of the CS contained a 3-prime Black Hole Quencher 2 (BHQ2) that would, after annealing, be in close proximity to the 5-prime cyanine-5 (Cy5) linked to the Trp aptamer.

The aptamer was hybridized with an excess of each CS in separate reactions, after which Trp was titrated to the solution (**Figure 2C and 2D**). The aptamer to which no CS was added displayed as expected the highest fluorescence, which was unaffected by the presence of Trp. The shortest CS, CS-9 (complementary strand of nine base pair length), lowered the fluorescence by 50% in the absence of Trp, indicating that 50% of the aptamer was still unbound in solution, and 50% of the aptamer was bound. The length of the CS lowered the background systematically, consistent with the idea of making the equilibrium more favourable with increasing base pairing interactions. CS-15 reduced the background to zero, quenching all the Cy5 fluorescence and highlighting the efficiency of the quenching method. CS-15, however, created a duplex which was too stable to be affected by the presence of Trp, blocking the Trp binding site and rendering it useless as a biosensor. Finally, CS-10 showed a concentration dependence in titration curves (**Fig 2C and 2D**) and was chosen for further optimization. The equilibrium favours binding of CS-10 in the absence of Trp, while CS-10 gets readily displaced in favour of Trp when present.

The hybridization of CS with the aptamer is dependent on both the concentration and stoichiometry of the CS. The higher the concentration of CS-10, the higher the percentage of bound species, lowering the background fluorescence. The effectiveness of increasing concentration saturated around 0.5 µM of the aptamer:CS10 complex, which was chosen for all further experiments (**Supplementary Figure 2A**). The background can be further reduced by increasing the equivalents of CS-10 to aptamer in solution. We found little additional benefit above a 2.5-fold excess of the CS (0.5 µM:1.25 µM Trp aptamer:CS10) after which the background signal seemed to stabilize (**Supplementary Figure 2B**). Under these conditions, we found the CS-10 to have the highest signal-to-noise ratio of all tested aptamer:CS complexes, which was ∼6-fold when 5 mM Trp was present (**Supplementary Figure 2C**). The *K*_sens_ of the Trp aptamer sensor was determined to be 3.3 mM, which is an order of magnitude weaker than the *K*_d_ of the original aptamer for Trp (1.8 µM) (**Supplementary Figure 2D**). The drop in *K*_sens_ compared to the *K*_d_ of the original aptamer was also observed in the similarly constructed biosensor for L-tyrosinamide.^18,23^ While the *K*_sens_ could be further improved by increasing the affinity of the aptamer for Trp, mM production of Trp in a droplet is already within reach considering the activity and expression levels of TrpB. Indeed, the expected concentrations of enzyme in a droplet from a single cell is estimated to range between 1 and 10 µM of enzyme, ^24^ so that no more than 1000 turnovers are necessary to create the mM quantities of Trp – well within reach of evolved TrpB variants.^20^

### Specificity of the Trp biosensor and compatibility with TrpB in plates and droplets

The hybridized Trp biosensor (Trp aptamer (0.5 µM) - CS10 (1.25 µM)) was combined with 5 mM of Trp, D-tryptophan or similar amino acids (**Figure 3A**). The Trp biosensor is specific for Trp and did not de-hybridize when similar amino acids such as L-phenylalanine were present. The aptamer sensor (much like the aptamer) is, however, sensitive to D-tryptophan. This is not a problem for TrpB evolution using L-serine as substrate. Although enantioselective aptamers have been evolved with SELEX, enantioselectivity was not a target criterion when the Trp aptamer was matured. The slight decrease in signal for D-tryptophan (as a result of slightly lower affinity) is likely to be reversed whenever the unnatural L-DNA duplex of the Trp sensor is hybridized due to reciprocal chiral substrate selectivity, and could become useful when selecting for D-tryptophan products.^19^

**Figure 3.**
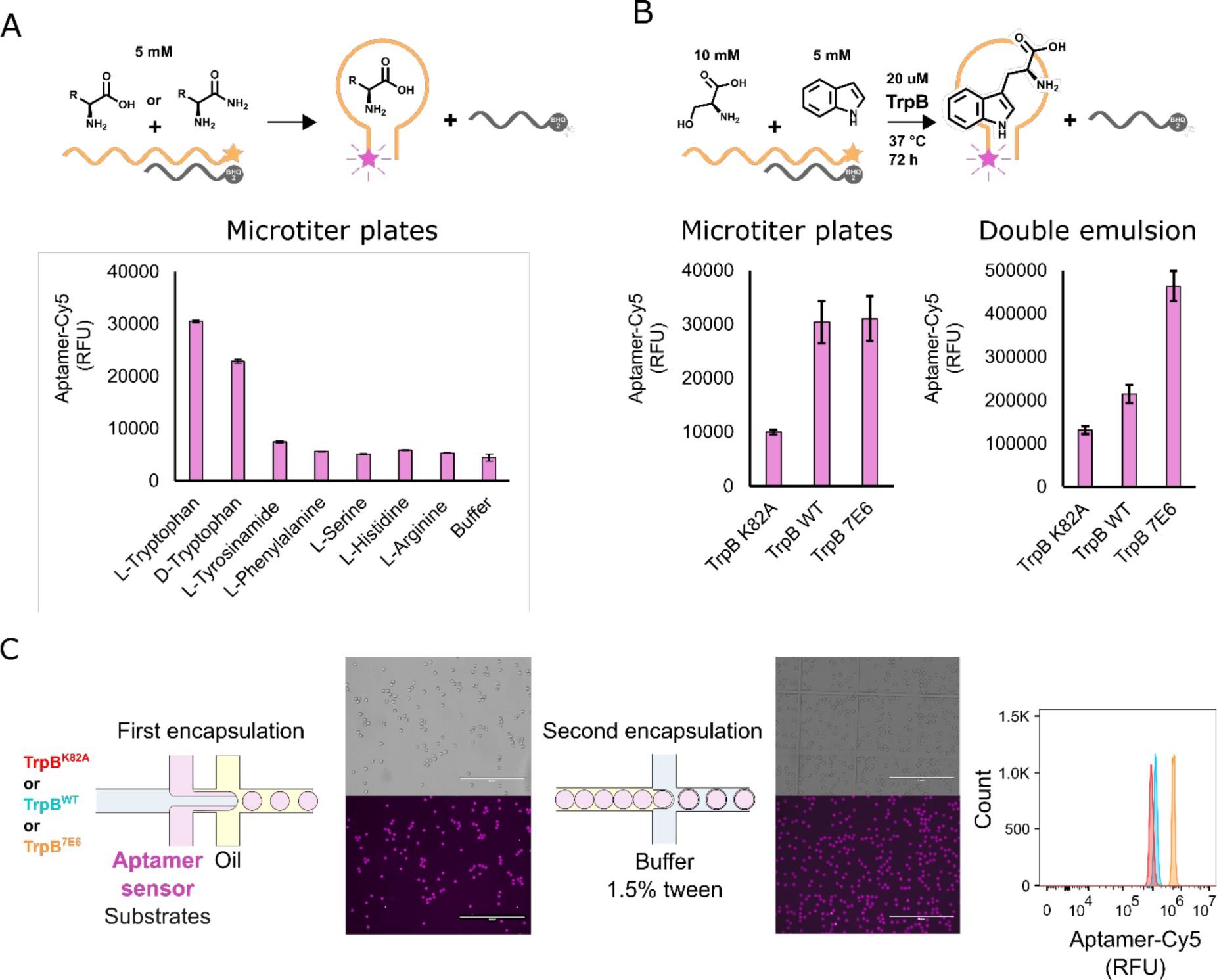
Specificity and compatibility of the Trp biosensor for Trp and Trp catalysis. (A) The Trp biosensor was hybridized and combined with 5 mM of each of the substrates shown in separate reactions, measuring the increase in fluorescence compared to when only TAB was added. (B). 10 mM Ser, 5 mM indole, 20 µM purified TrpB, and the aptamer sensor were combined and left to react at 37 °C for 72 hours to assess the stability of the sensor. After 72 hours, the mixtures were cooled down to RT and analyzed either by UV-Vis (for wells containing the solution), or via flow cytometry (for double emulsions containing the solution). (C) Encapsulation of 20 µM TrpB, with 10 mM Ser and 5 mM indole with the Trp biosensor. The droplets were incubated for 72 hours before being encapsulated again and prepared for flow cytometry. All raw data for the double emulsions are shown in **Supplementary Figure 4**.

Next, the compatibility of the Trp biosensor with the enzymatic production of Trp was probed. Three variants of TrpB were purified: TrpB^K82A^, TrpB^WT^ and TrpB^7E6^. TrpB^K82A^ functions as a catalytically inactive control, as the K82 residue connects the pyridoxal phosphate cofactor to the protein backbone and was mutated to an alanine. TrpB^7E6^ functions as a high activity comparison, which contains nine mutations compared to wildtype, accumulated over seven rounds of directed evolution/rational engineering, gradually increasing the expression levels, TTNs and substrate scope of TrpB.^25^ First, both TrpB^WT^ and TrpB^7E6^ were assessed for their compatibility with Trp aptamer buffer (TAB; 10 mM Na_2_HPO_4_, 2 mM KH_2_PO_4_, 2.7 mM KCL, 5 mM MgCl_2_, 500 mM NaCL, pH 7.4), set up to stabilize the secondary structure of the aptamer by high ionic strength and high magnesium concentrations (**Supplementary Figure 3**). The biosensor is sensitive to changes in the buffer components (requiring 1-10 mM Mg^2+^ and 100-1000 mM NaCl, **Supplementary Figure 4**), suggesting that reaction buffer conditions should be tailored for each new aptamer sensor.^21^ In TAB, both variants complete turnover of 5 mM of indole with Ser to produce 5 mM Trp (chosen as a benchmark as it is well above the K_sens_ of the biosensor, 3.3 **Supplementary Figure 2D**) in a matter of ∼6 hours. For comparison, in a KP_i_ buffer (in which TrpB^7E6^ was evolved)^25^, the enzymes require <1 hour for complete turnover.

The purified proteins were combined with Ser, indole, and the Trp sensor in microtiter plate format and left to react for ∼72 hours to probe the stability of the sensor under reaction conditions (**Figure 3B**). In the presence of TrpB^WT^ or TrpB^7E6^ a clear increase in fluorescence suggests that production of Trp can be monitored, compared to a ∼4.5-fold smaller fluorescence change in TrpB^K82A^ containing wells that does not produced Trp. The TrpB^K82A^ control highlights the specificity of the aptamer for Trp over its substrates and any tryptophan on the surface of TrpB, reducing the dynamic range compared to buffer only conditions only marginally from ∼6 fold to ∼4.5 fold.

Using the same concentrations, TrpB variants, substrates and Trp sensor were combined in microfluidic droplets and left to incubate for 72 hours (**Figure 3C**). Flow cytometric analysis of the separate populations of double emulsion droplets is shown in **Supplementary Figure 5**. Again, the Cy5 fluorescence of the sensor increased in droplets where active TrpB was present compared to TrpB^K82A^, which for TrpB^7E6^ mimicked the dynamic range observed in microtiter plate-based format. The TrpB^WT^ containing droplets deviated from the plate-based results, which, although more active than TrpB^K82A^, did not completely turnover indole (**Figure 3B**). This could be explained by the mandatory addition of surfactant Tween-20 to stabilize double emulsions, or the presence of an oil interface when performing the catalysis in droplets. Nevertheless, the aptamer is both selective, and compatible with enzymatic catalysis of Trp by TrpB variants.

### Enrichment of active TrpB in plate-based and droplet-based format with the Trp biosensor

The Trp biosensor is compatible with purified TrpB, but for directed evolution, TrpB needs to be expressed in single wells or single cells to screen a library of enzyme variants. *E. coli* contains endogenous tryptophanase (TnaA), which can degrade up to 5 mM of exogenous Trp to produce indole.^26^ To facilitate screening with *E. coli*, a tryptophanase-deficient cell strain was used, *E. coli* DE3 (C43) ΔTnaA, previously engineered in the Bernardt group.^27^ The removal of TnaA was shown not to affect the growth rate of *E. coli* and was, therefore, well suited for directed evolution experiments.^27^

First, we probed the Trp biosensor in a microtiter plate-based assay. TrpB^7E6^ and negative control (plasmid without insert) were transformed and expressed in a 1 mL culture of *E. coli* DE3 (C43) ΔTnaA. Next, the fluorescence was measured after sequential addition of both Ser, indole and the aptamer sensor (**Supplementary Figure 6**). Each of the wells containing cells expressing TrpB^7E6^ were of higher fluorescence than those containing cells expressing no TrpB. As such, we were confident that the Trp biosensor is able to localize TrpB activity expressed from a 1-mL culture.

Given the potential of non-canonical amino acids as precursors to pharmaceuticals, we probed whether the Trp biosensor could be used to screen for value-adding, Trp derivatives (**Supplementary Figure 7**). We found that the biosensor is versatile enough to distinguish both 5-fluoro-L-tryptophan and 5-methoxy-L-tryptophan from the respective precursors. Given the more limited solubility of 5-methoxyindole, we used 5-fluoroindole as the substrate for testing the enrichment of TrpB^7E6^ variant in droplets.

Both positive and negative controls were transformed into *E. coli* DE3 (C43) ΔTnaA again, this time aiming to encapsulate single cells in droplets with 5-fluoroindole as the substrate. Before encapsulation, cells transformed with an empty vector control were mixed with TrpB^7E6^ transformed cells in a 100:1 ratio. After incubation, the droplets were sorted via FACS, and the genotype was recovered from both the sorted and unsorted fractions (**Figure 4**). The recovered genotype was transformed into *E. coli* DE3 (C43) ΔTnaA, picking individual colonies for a plate-based rescreen using the aptamer sensor (**Figure 4**). As predicted, the unsorted fraction contained only 2% of TrpB^7E6^, closely matching the intended ratio. The sorted fractions, containing the top 0.7% and 0.1% fluorescent droplets, however, contained 36% and 45% TrpB^7E6^ variants after sorting (and confirmed by sequencing), which constitutes a ∼40x enrichment of the originally intended ratio of active and empty vector control. Having shown enrichment for active TrpB in both plate-based and droplet-based formats, we started evolving TrpB.

**Figure 4.**
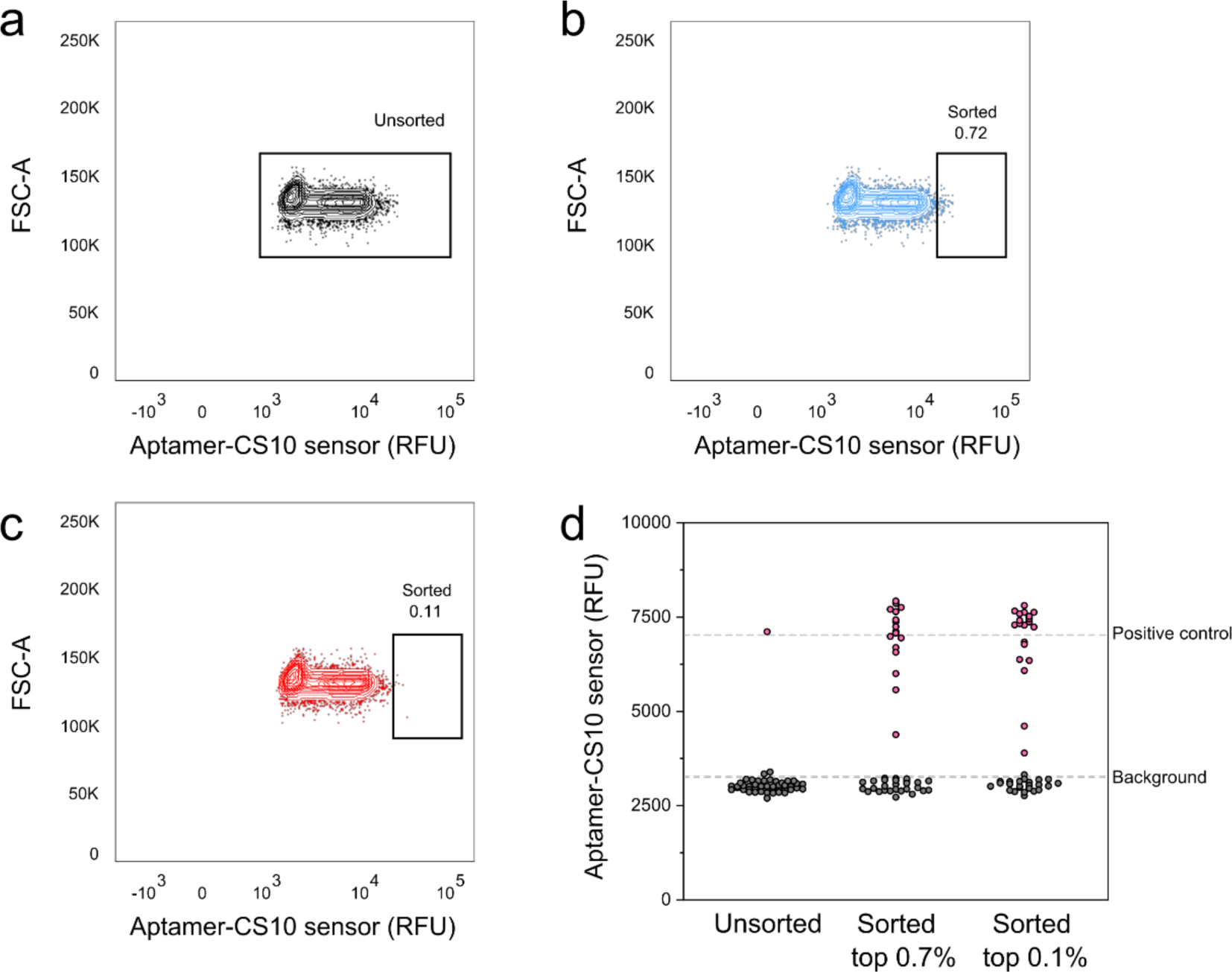
Enrichment of TrpB7E6 in double emulsions. Cells expressing either TrpB^7E6^ or the vector without insert were mixed in a 1:100 ratio, and encapsulated with 5-fluoroindole, Ser and lysis reagents. After 72 hours of incubation, the droplets were encapsulated again and sorted via FACS. The recovered genotype from the unsorted fractions (a), top 0.7% (b) and top 0.1% (c) was transformed and rescreened in plates for 5-fluoro-_L_-tryptophan production using the aptamer sensor. The fluorescent intensity of all wells is shown in d. Additionally, the plates were sequenced to confirm the high fluorescent wells to contain TrpB^7E6^, which are denoted as purple-coloured dots, where no false negatives were observed.

### Evolution of TrpB

The first round of evolution from the starting point, TrpB^WT^, was done in plates using the aptamer sensor, focussing on finding an activating and/or strongly expressing variant that would benefit the sensitivity of screening in droplets. An error-prone library was prepared with 2.2 ± 1.4 mutations per gene. Out of ∼200 variants, the top variant, TrpB^B10^ was isolated, sequenced and expressed. The variant contained one mutation, F176L, which increased expression levels of TrpB from 50 mg/L to 135 mg/L, while additionally moderately increasing the number of TTNs and catalytic efficiency **(Table 1; Supplementary Figure 10)**. Combined, we deemed the TrpB^B10^ variant to be well-suited as a template for directed evolution in the droplet-based screen.

**Table 1.**
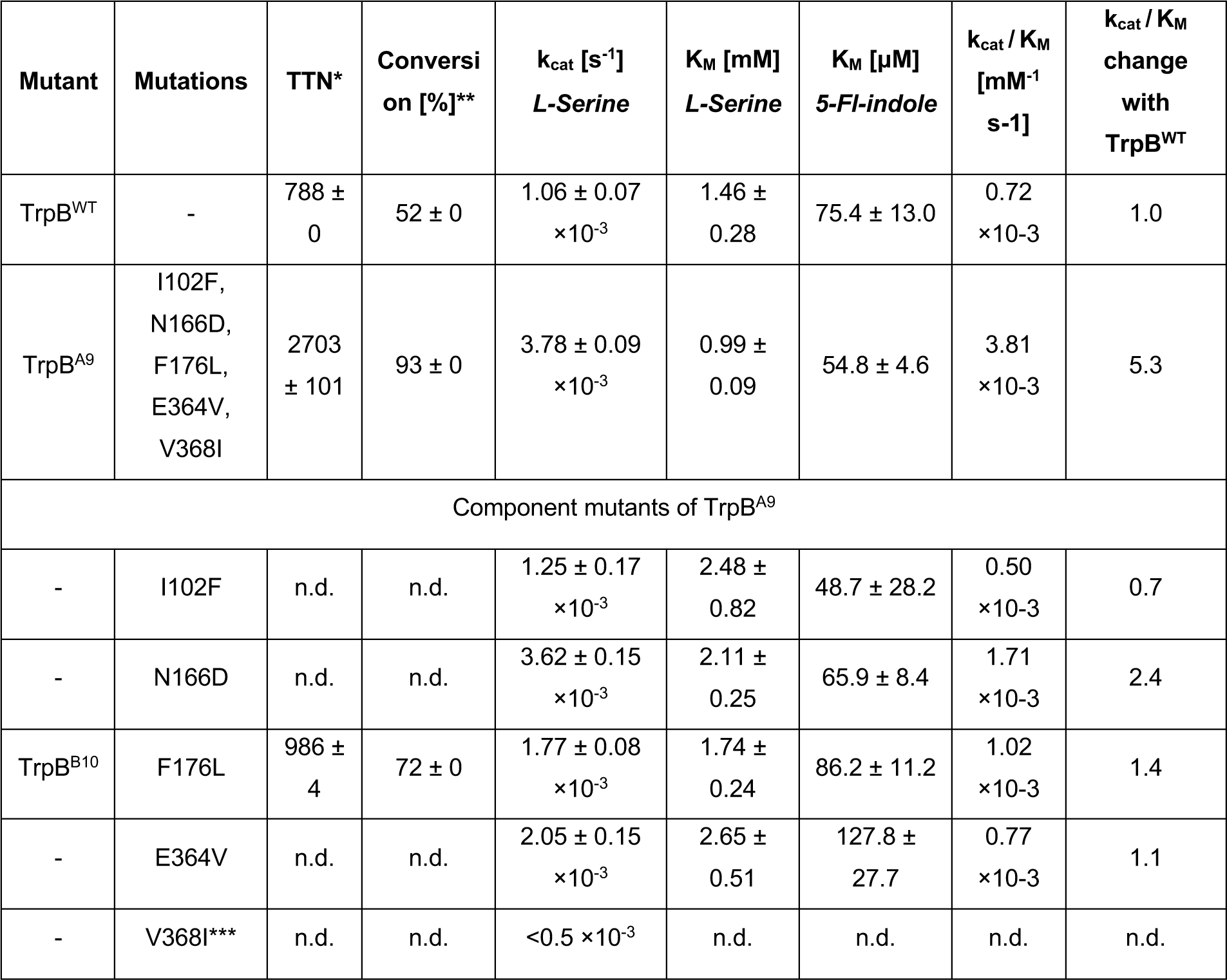
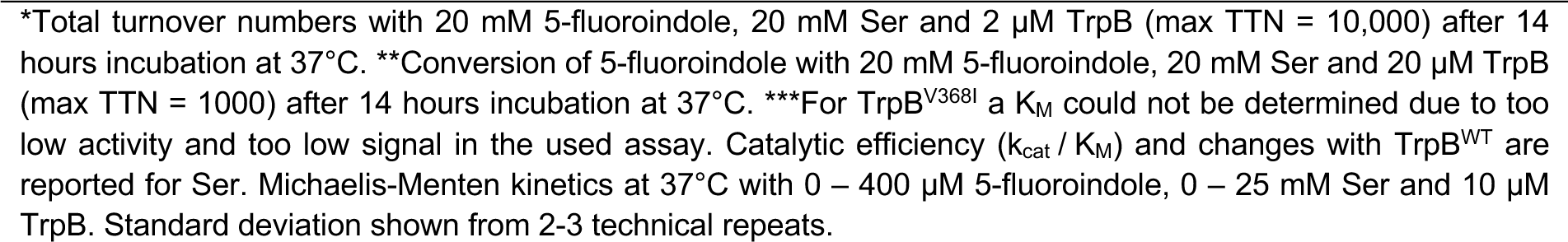
Characterization of evolved TrpB variants.

| Mutant | Mutations | TTN* | Conversion [%]** | $k_{cat} [\text{s}^{-1}]$<br>*L-Serine* | $K_M [\text{mM}]$<br>*L-Serine* | $K_M [\mu\text{M}]$<br>*5-Fl-indole* | $k_{cat} / K_M$<br>[$\text{mM}^{-1} \text{s}^{-1}$] | $k_{cat} / K_M$<br>change<br>with<br>TrpB<sup>WT</sup> |
| --- | --- | --- | --- | --- | --- | --- | --- | --- |
| TrpB<sup>WT</sup> | - | 788 ± 0 | 52 ± 0 | $1.06 \pm 0.07 \times 10^{-3}$ | $1.46 \pm 0.28$ | $75.4 \pm 13.0$ | $0.72 \times 10^{-3}$ | 1.0 |
| TrpB<sup>A9</sup> | I102F,<br>N166D,<br>F176L,<br>E364V,<br>V368I | 2703 ± 101 | 93 ± 0 | $3.78 \pm 0.09 \times 10^{-3}$ | $0.99 \pm 0.09$ | $54.8 \pm 4.6$ | $3.81 \times 10^{-3}$ | 5.3 |
| Component mutants of TrpB<sup>A9</sup> |  |  |  |  |  |  |  |  |
| - | I102F | n.d. | n.d. | $1.25 \pm 0.17 \times 10^{-3}$ | $2.48 \pm 0.82$ | $48.7 \pm 28.2$ | $0.50 \times 10^{-3}$ | 0.7 |
| - | N166D | n.d. | n.d. | $3.62 \pm 0.15 \times 10^{-3}$ | $2.11 \pm 0.25$ | $65.9 \pm 8.4$ | $1.71 \times 10^{-3}$ | 2.4 |
| TrpB<sup>B10</sup> | F176L | 986 ± 4 | 72 ± 0 | $1.77 \pm 0.08 \times 10^{-3}$ | $1.74 \pm 0.24$ | $86.2 \pm 11.2$ | $1.02 \times 10^{-3}$ | 1.4 |
| - | E364V | n.d. | n.d. | $2.05 \pm 0.15 \times 10^{-3}$ | $2.65 \pm 0.51$ | $127.8 \pm 27.7$ | $0.77 \times 10^{-3}$ | 1.1 |
| - | V368I*** | n.d. | n.d. | $<0.5 \times 10^{-3}$ | n.d. | n.d. | n.d. | n.d. |

\*Total turnover numbers with 20 mM 5-fluoroindole, 20 mM Ser and 2 μM TrpB (max TTN = 10,000) after 14 hours incubation at 37°C. \*\*Conversion of 5-fluoroindole with 20 mM 5-fluoroindole, 20 mM Ser and 20 μM TrpB (max TTN = 1000) after 14 hours incubation at 37°C. \*\*\*For TrpB<sup>V368I</sup> a $K_M$ could not be determined due to too low activity and too low signal in the used assay. Catalytic efficiency ( $k_{cat} / K_M$ ) and changes with TrpB<sup>WT</sup> are reported for Ser. Michaelis-Menten kinetics at 37°C with 0 – 400 μM 5-fluoroindole, 0 – 25 mM Ser and 10 μM TrpB. Standard deviation shown from 2-3 technical repeats.

An error-prone library of TrpB^B10^ was prepared with 4.5 ± 2.5 mutations per gene. For the droplet screen, the mutation frequency was increased and this increase in screening throughput was thought to counteract the expected decrease in expected functional variants.^28^ With this high mutational rate, we set out to enrich for variants that combine 4-5 new, activating or neutral mutations per round. Single cells harbouring unique TrpB variants were encapsulated with 5-fluoroindole, Ser, and lysis reagents, incubated, encapsulated again and sorted (**Supplementary Figure 8)**. TrpB^B10^ was expected to turn over 5 mM of product for a maximal signal during incubation (**Supplementary Figure 3**). The top 5.7% of fluorescent double emulsion droplets were sorted from a total of 234,000 droplets, and the genotype was recovered and rescreened in plate-based format. We observed an average activity increase of 32% in the sorted fraction, and a 44% increase in activity for the top 25 percent of rescreened wells (*M* (3939), *Q3* (5111) compared to that of the unsorted library (*M* (2988)*, Q3* (3548) (**Supplementary Figure 9**).

From the ∼13,000 sorting events, 400 variants were rescreened for 5-fluoroindole conversion in plates using the aptamer sensor. The seven variants with the highest activity were purified and combined with 5-fluoroindole and Ser, testing the TTN (0.01% catalyst loading, max TTN = 10,000) and conversion (0.1% catalyst loading, max TTN = 1000) on HPLC. All variants yielded improvements in TTN, conversion, and/or expression yields **(Supplementary Figure 10)**. The variant with the highest TTN’s, TrpB^A9^, increased the catalytic efficiency by more than 5-fold from 0.72 ×10^-3^ to 3.81 ×10^-3^ mM^-1^ s^-1^ compared to TrpB^WT^, due to improvements in the K_m_ for both Ser and indole, and an increased k_cat_ **(Table 1)**. The variant TrpB^A9^ carries additional four mutations (I102F, N166D, E364V, V368I) to the F176L mutation of its parent TrpB^B10^. Interestingly, in terms of *k*_cat_, the mutations are almost perfectly additive, i.e. the *k*_cat_ of the combined mutant TrpB^A9^ closely resembles the expected fold change from the *k*_cat_ changes of its component mutations. When multiplying the *k*_cat_ ratios of TrpB WT and each component mutant: F176 (1.67), I102F (1.18), N166D (3.42), E364V (1.94), V368I (0.26) with the *k*_cat_ of TrpB WT (1.06 ×10^-3^), the expected *k*_cat_ (3.70×10^-3^ s^-1^) closely resembles the observed *k*_cat_ of TrpB^A9^ (3.78 ×10^-3^ s^-1^) **(Table 1)**. (Because the *K*M of TrpB V368I could not be measured with confidence, similar epistatic analysis for catalytic efficiency was not possible.) These observations suggest that operating in a high mutagenesis regime, beneficial and synergistic combinations of residue mutations are selected.

Additionally, TrpB^A9^ improved activity by ∼2.5 fold on the indole derivatives 5-methyl, 6-methyl, 6-fluoro, 5-fluoro, 5-hydroxy and 5-methoxy indole compared to TrpB^WT^, to produce the corresponding tryptophan derivatives **(Table 2)**.

**Table 2.**
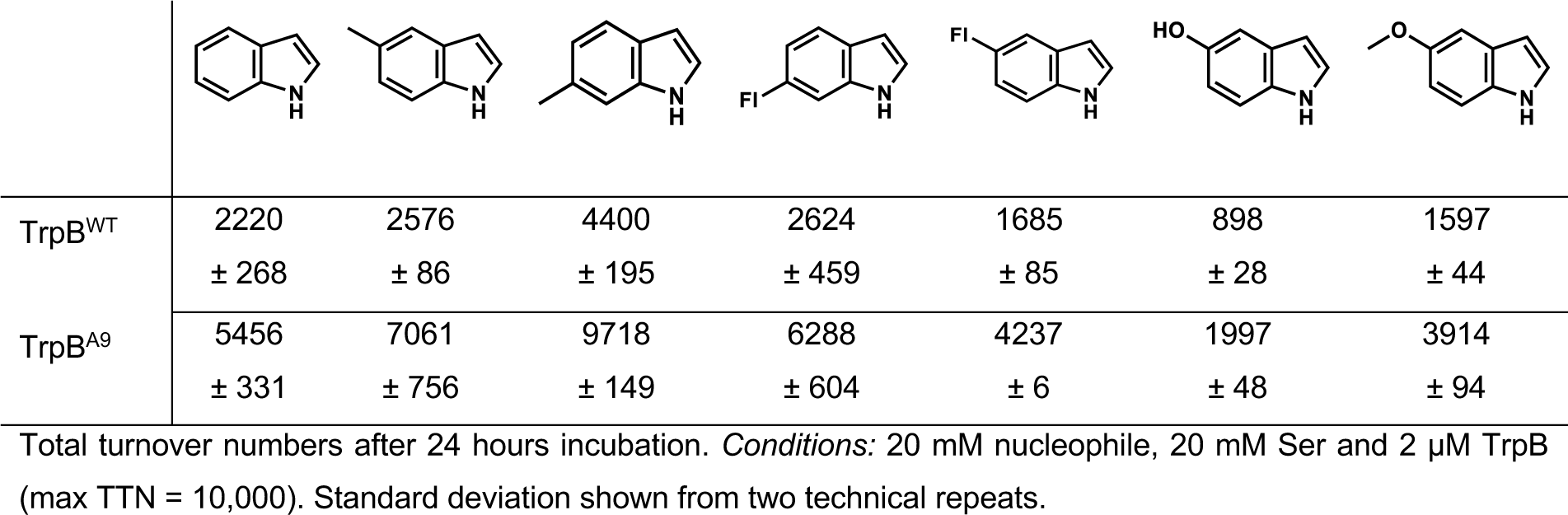
Substrate scope of TrpB^A9^.

|  |  |  |  |  |  |  |  |
| --- | --- | --- | --- | --- | --- | --- | --- |
| TrpB<sup>WT</sup> | 2220<br>± 268 | 2576<br>± 86 | 4400<br>± 195 | 2624<br>± 459 | 1685<br>± 85 | 898<br>± 28 | 1597<br>± 44 |
| TrpB<sup>A9</sup> | 5456<br>± 331 | 7061<br>± 756 | 9718<br>± 149 | 6288<br>± 604 | 4237<br>± 6 | 1997<br>± 48 | 3914<br>± 94 |

Total turnover numbers after 24 hours incubation. *Conditions:* 20 mM nucleophile, 20 mM Ser and 2 μM TrpB (max TTN = 10,000). Standard deviation shown from two technical repeats.

## Discussion

Combining microfluidics with the versatility of SELEX to create binders for small molecules enables screening for improved enzymatic production of any product at ultra-high throughput. A myriad of aptamers for small molecules have been evolved through SELEX, highlighting its versatility ^14,15^. The most important criterion of an aptamer used in enzyme screening is differential recognition of the product of interest over the substrate(s) that it originates from. To guarantee this specificity, negative selection rounds can be employed to counter-select for substrate binding, as exemplified by an aptamer for L-arginine with a 10,000:1 preference for the L-over the D-enantiomer.^29^ Here, we chose to use an existing aptamer for Trp,^21^ which showed minimal binding of the substrates from which Trp is assembled: indole and Ser. Tryptophan-producing enzymes are interesting targets for directed evolution as derivatives of Tryptophan – non-canonical amino acids – are found in 12% of the 200 top-grossing pharmaceuticals, ^30,31^ and are notoriously difficult to synthesize using conventional synthetic chemistry.

After a specific aptamer for the product of interest is obtained, the aptamer is engineeringed into a sensor. The Heemstra group has developed a straightforward approach, requiring the synthesis of <10 complementary strands with a quencher, complexing it and titrating the product of interest to check for signal-to-noise ratio. Here, we applied the same strategy initially developed for a L-tyrosinamide aptamer and applied it to a Trp aptamer. We found the strategy to be generalizable: our optimal CS was only one base pair longer than the CS for the L-tyrosinamide aptamer, whereas the optimal stoichiometry and concentration of aptamer to CS was found to be identical. The resulting pipeline simplifies the screening approach compared to RNA-aptamers-in-droplets (RAPID) screening approach.^17^ First, for DNA-aptamer sensors, there is no need for in silico optimization to predict the best aptamer-Spinach hybrids. Second, RNA is inherently less stable and requires IVT and subsequent purification to be produced. DNA aptamers can be readily supplied by commercial manufacturers as fluorescent single-strand oligos and annealed with complementary quenching strands to create a stable duplex. Lastly, the use of DMHBI, which complexes with the folded Spinach aptamer for fluorescence, is not required as the aptamer itself is already labelled with fluorescent dye.

Whereas the Heemstra group used the aptamer sensor primarily to differentiate between L-tyrosinamide and D-tyrosinamide, here we applied the concept to screen for improved enzyme variants in droplets. To make the screening approach more available to the wider user base, the concept was developed in a double emulsion droplet format compatible with a commercial FACS instrument.^32^ (Double emulsion droplet sorting may require some optimization, which is well summarized in the supplementary information of Brower et al.^33^) We found, with good reproducibility, that the majority of our droplets (80%) were individual double emulsions droplets so that droplet fusions and oil droplets were of minimal concern. Furthermore, FACS allows for gating around the correct droplet population so that a more homogeneous sample is sorted, which provides an advantage over fluorescent sorting of droplets on chips where larger droplets as a result of fusion cannot easily be practically excluded before sorting.

As such, the setup provides experimental risk mitigation, as the double emulsions can be prepared using a setup with three pumps and a camera, after which the double emulsions can be stored in buffer in standard tubes and sorted on FACS. Thus allowing screening of enzyme variants with little reagent consumption, without the need to invest in specialized on-chip sorting devices. Other non-labelled droplet sorting techniques based on RAMAN or Mass Spectrometry (MS) function independent of aptamer maturation, and are not limited by the buffer sensitivity of aptamer folding.^8,34,35^. Although RAMAN and MS sorting do not require SELEX, and are therefore intrinsically more flexible, both methods require advanced set-ups and are currently only possible at low sorting frequencies (sorting 1-10 Hz, i.e. < 9×10^5^ droplets per day), whereas FACS can sort 10^4^ – 10^5^ droplets per second and is therefore only limited by droplet formation (creating 10^3^-10^4^ droplets per second). The aptamer sensor has lower sensitivity compared to MADS (mid µM). The *k*_sens_ of the tryptophan sensor was 3.3 mM with a ∼6 fold dynamic range across experiments **(Supplementary Figure 2D)**, with a lower limit of detection of 1 mM (**Supplementary Figure 2C**). Due to the solubility of Trp, the dynamic range is limited at 5 mM, where the fluorescent signal was shown to saturate **(Supplementary Figure 2C)**. Increasing the sensitivity can be achieved by selecting higher affinity aptamers during in the initial SELEX, which, as a result, can be complexed with a longer CS, so that background fluorescence is reduced and and lthe *k*_sens_ lowered.

The Trp aptamer droplet-based uHTS approach was utilized to screen a library with a high mutational rate, in departure from the widespread practise to explore sequence space ‘*one amino acid at a time’*.^36,37^ Increasing the number of mutations per gene lowers the chance of finding active variants and, therefore, requires increased throughput to find variants of interest.^28^ We obtained a TrpB variant which accumulated 4 mutations in a single round, making it 5-fold more active than WT TrpB at 37°C. The improvement is similar to what was achieved in previous low throughput mutagenesis studies of TrpB from the thermophile *Pyrococcus furiosus*. The first study, evolved TrpB for Trp over 3 rounds, which resulted in 83-fold increase in catalytic efficiency at 75°C (averaging 2 mutations per round, and ∼28-fold activity increase per round).^38^ Next, TrpB^4D11^ was evolved for β-methyl-Trp over 3 rounds, which resulted in a 7-fold improvement in β-methyl-Trp production over 2 rounds (averaging 1 new mutation and 3.5-fold increased activity per round).^39^ After introduction of a rationally designed active site mutation, TrpB^L161A^ was evolved for β-ethyl-Trp over 3 rounds, which resulted in a 6-fold increase in TTN (averaging 1 mutation and 2-fold activity increase per round).^25^ Recently, the continuous directed evolution platform OrthoRep was utilized to evolve TrpB from *Thermotoga maritima* (*Tm*TrpB) in ultra-high throughput.^40^ Over 13-20 passages, or 40 days of continuous evolution, an average of 10 mutations/gene were introduced, increasing catalytic activity of *Tm*TrpB by 22-fold at 37°C. The Trp-producing gene of the screening host was knocked out, so that the proliferation of cells was directly dependent on the production of Trp by *Tm*TrpB. While uncovering a plethora of new mutations (∼200) in *Tm*TrpB that accelerates the evolution of promiscuous activities, it is unlikely that the strategy employed is adaptable to products which are not essential for cell growth. Even though our dynamic range is small (and practically capped at production of 5 mM of Trp) we envision increasing the stringency of selection by decreasing the incubation time so that increases in *k*_cat_ are made evident, over multi rounds of mutagenesis with high mutational load.

Interestingly, the most active component mutation in TrpB^A9^, N166D had been localized both in the first low-throughput evolution of TrpB, and in the continuous evolution of *Tm*TrpB (corresponding to N167D).^38,40^ Additionally, the positions of component mutation F176L and E364V were also found as beneficial sites in *Tm*TrpB (corresponding to mutations L177Q, and K361E respectively).^40^ Most of the mutations uncovered here (I102F, N166D, and F176L) are found in the COMM domain of TrpB.^41^ Mutations in this domain were shown to increase the activity of TrpB by mimicking allosteric activity by TrpA – resulting in a closed conformation of TrpB, changing the rate-limiting step of the catalytic cycle.^42^ Indeed, the mutations uncovered are likely to act through a general mechanism outside of the active site, as they do not only improve activity on 5-fluoroindole, but increase activities for multiple substrates of TrpB.

Considering the abundance of negative epistasis (estimated to affect ∼50% of pairwise interactions)^43^, it is interesting to observe that the component mutations were almost perfectly additive. Although screening with a high mutational rate does not preclude observing activating variants with hitchhiking mutations which negatively affect overall activity, it can select for combinations of activity-decreasing mutations which together form a more active enzyme through reciprocal sign epistasis. The synergy of multiple residues cannot be addressed through low-throughput mutagenesis e.g. by assessments of mutability^44–46^ (which will evaluate single positions iteratively by mutational scanning). Simultaneous mutagenesis in multiple positions is therefore likely to give rise to residue combinations that break new ground and can be visualised as novel fitness peaks, even in a rugged landscape.^47^

Taken together, the aptamer strategy employed here opens up possibilities to find fitness peaks which are often clouded through negative epistasis in faraway regions of sequence space. More importantly, we hope that the generalizability of DNA aptamers and double emulsions facilitate high throughput screening at a scale that paves the way for new strategies in directed evolution campaigns, with minimal capital investment and low consumables expenditure due to minaturised assay volumes.

## Acknowledgements

We thank Joana Cerveira of the Department of Pathology for her help with flow cytometric analysis. This work was funded by the Horizon 2020 programme of the European Commission. R.A.S. was supported by a studentship from the EU ITN MMBIO and F. E. H. N. the EU ITN Oligomed. T.S.K. was supported by EU H2020 Marie Skłodowska-Curie Individual Fellowship (MSCA-IF 750772) and F.H. is an ERC Advanced Investigator (695669).

## Author contributions

Conceptualization, R.A.S.; Methodology, R.A.S., Y.W. and T.S.K.; Investigation R.A.S, Y.W. F.E.H.N. and M.H.; Data Curation, R.A.S, F.E.H.N. and M.H.; Writing – Original Draft, R.A.S.; Writing – Review & Editing, R.A.S., F.E.H.N., T.S.K. and F.H.; Visualization, R.A.S.; Supervision, F.H.; Funding Acquisition, F.H.

## Materials and Methods

### Cloning

TrpB^WT^ and TrpB^7E6^ from *Pyrococcus furiosus* were ordered as genestrings, and next were subcloned to pRSF, a kind gift from J. D. Schnettler. TrpB variants were amplified using Q5 2x MM (NEB): 98°C for 2 minutes, 98 °C for 10 seconds, 67 °C for 15 seconds, 72°C for 1 minute, and final extension at 72°C for 5 minutes. The accepter plasmid, pRSF, was amplified with Q5 2x MM: 98°C for 2 minutes, 98 °C for 10 seconds, 68 °C for 15 seconds, 72°C for 3 minutes, and final extension at 72°C for 5 minutes. The TrpB genes were assembled in pRSF using Gibson assembly. TrpB^K82A^ was prepared using site-directed mutagenesis (NEB), according to protocol. Protein and DNA sequences in **Supplementary Table 1**.

Error-prone libraries were prepared using GeneMorph II Random Mutagenesis Kit (Agilent). The fragments were purified from agarose gels over silica using agarose dissolving buffer (Zymoclean Gel DNA Recovery, Zymo Research). The library fragments were cloned into pRSF accepter vector described above. The Gibson fragment was purified over silica columns and transformed to *E. cloni* 10F ELITE Electrocompetent cells (Lucigen), and plated on two 140 mm Petri dishes containing LBkan-agar. The next day, the colonies were scraped, and the plasmid was isolated using Genejet plasmid miniprep kit (Qiagen). This purified plasmid stock was used for transformation BL21 (DE3) competent *E. coli* (NEB, 2527) and *E. coli* C43 DE3 ΔTnaA for plate screening experiments, and to *E. coli* C43 DE3 ΔTnaA for microfluidic experiments.

### Protein expression and purification

Purified pRSF_TrpB, pRSF_TrpB^7E6^ and pRSF_TrpB^K82A^ plasmids from a single colony were transformed to *E. coli* BL21 (DE3) and plated on LB-Agar with the appropriate selection marker(s). The next day, the colonies were scraped and used to inoculate 500 mL of LB and grown until an OD600 of ∼0.4-0.8. Cells were induced with 1 mM IPTG. Expression was done overnight at 25°C, 200 rpm.

Cells were harvested the next day and washed with Binding Buffer (25 mM KPi, 100 mM NaCl, 5 mM Imidazole, pH 8.0). 15 mL Lysis Buffer (Binding buffer, 1x Bugbuster, 1 mg/mL hen egg white lysozyme (HEWL), 200 µM PLP, 125 U/mL Benzonase, pH 8.0) was added, manually resuspended and incubated at room temperature for 30 minutes. Cell debris was palleted (15,000 rpm, 30 minutes), and the supernatant was applied to a gravity his-tag column (CV = 2 mL). The resin was washed with 3 CV Wash buffer (25 mM KPi, 100 mM NaCl, 30 mM Imidazole, pH 8.0), before being eluted with 5 mL Elution buffer (25 mM KPi, 100 mM NaCl, 500 mM Imidazole, pH 8.0). The yellow fraction was collected. The eluted product was dialyzed with PD-10 columns according to manufacturer’s protocol (GE Healthcare) to Trp Aptamer buffer (TAB, (2 mM KH2PO4, 10 mM Na2HPO4, 2.7 mM KCl, 5 mM MgCl2, 500 mM NaCl, pH 7.4)) for all experiments, and 50 mM KPi as a control when optimizing buffers.

Proteins were flash-frozen in liquid nitrogen, aliquoted and stored at −80°C. Thawed aliquots were only used once, and discarded the same day.

### Preparation of tryptophan sensor

Tryptophan aptamer and complementary strands were diluted in Trp aptamer buffer (TAB (10 mM Na2HPO4, 2 mM KH2PO4, 2.7 mM KCL, 5 mM MgCl2, 500 mM NaCL, pH 7.4)). The aptamer sensor was prepared by generating a 2x solution of 1 µM aptamer with 2.5 µM CS-10. For microfluidic experiments, a 4x solution of 2 µM aptamer with 5 µM CS-10 was prepared. The mixtures were incubated for 10 minutes at 90°C, before being cooled down rapidly in a thermocycler to 4°C after which the sensor was immediately placed on ice for 10 minutes. The sensor was placed at room temperature for 10 minutes or longer before use, and never kept overnight.

### UV-Vis Spectroscopy

The aptamer sensor was combined 1:1 either with purified chemicals, or with TrpB reaction mixtures, in a final volume of 80 µL in low binding microtiter plates (Corning, 3881). The increase in fluorescence was measured over 90 minutes (25°C) on a Tecan infinite 200Pro using a 650 nM excitation wavelength, and measuring the 700 nm emission wavelength. The bandwidth of excitation/emission was 9/20 nm respectively, with 25 flashes per measurement. The increase in fluorescence was typically saturated after 90 minutes incubation, and taken as the end-point measurement reported.

### Catalysis with TrpB

All analytical reactions were performed in 2 mL HPLC glass vials. A solution consisting typically of 20 mM L-serine (from 0.5 M stock in ddH2O), 20 mM Indole (from 0.5 M stock in DMSO, 4% DMSO final), 200 µM PLP (from 10 mM stock in ddH2O) was combined with purified TrpB (in TAB). Reactions were incubated for 24 hours in a 37°C water bath.

#### Read-out with aptamer sensor

The mixture was cooled down to room temperature and combined 1:1 with the aptamer sensor before measuring for 90 minutes by UV-Vis spectroscopy as described above.

#### Read-out with HPLC

The mixture was quenched 1:1 with acetonitrile, and centrifuged at >14,000g for 10 minutes. The supernatant was analyzed by HPLC. HPLC was performed on an Agilent 1260 infinity II, equipped with a C-18 column (Mackerey-Nagel, 5 µm, ref 760.100.40) using acetonitrile (HPLC grade, Agilent) and ddH2O (0.1% (v/v) formic acid (FA)). The program was as follows: 0 min – 100% ddH2O (0.1% (v/v) FA) 0% acetonitrile. 3 min – 40% ddH2O (0.1% (v/v) FA) 60% acetonitrile. 5 min – 0% ddH2O (0.1% (v/v) FA) 100% acetonitrile. 6 min – 0% ddH2O (0.1% (v/v) FA) 100% acetonitrile. 7 min – 100% ddH2O (0.1% (v/v) FA) 0% acetonitrile All samples were analyzed at 277 nm, representing the isosbestic point between indole and tryptophan, allowing for estimation of yield by comparing the area of the substrate peak to the areas of both substrate and product peak combined.^48^

### Michaelis-Menten kinetics

Kinetic measurements of TrpB^wt^ and its mutants were performed by monitoring 5-fluoro-L-Trp formation in a plate reader (Thermo Scientific Varioskan Lux) over 20 min at 290 nm using *ΔE290 = 1.89 mM^−1^·cm^−1^*. Measurements were taken every 10 seconds and slopes were normalised on background absorbance changes. Initial rates were calculated using a L-tryptophan standard curve as proxy for 5-fluoro-tryptophan formation. All assays were conducted in a quartz 96-well plate at 37°C using 10 µM TrpB and 20 µM PLP in TAB with a final concentration of 1% DMSO. For serine kinetics, 0 - 25 mM L-serine and 200 µM 5-fluoroindole were used. For indole kinetics, 0 – 400 µM 5-fluoroindole and 25 mM L-serine were used. All measurements were performed in three technical replicates. Data was fitted to the Michaelis-Menten equation using Origin 2018 (Origin Lab).

### Plate screening

Individual *E. coli* BL21(DE3) or *E. coli* C43 DE3 ΔTnaA colonies containing either a variant for rescreening, or the TrpB^wt^ and TrpB^7E6^ controls were picked in 300 µL TBkan and grown overnight at 37°C, ∼900 rpm in a plate shaker. The next day, 30 µL was taken to inoculate 920 µL TBkan and grown for 3 hours at 37°C, ∼900 rpm. The plates were inoculated with 50 µL TBkan containing 20 mM IPTG (1 mM final) and grown overnight at 20°C. The next day, cells were palleted (4000 rpm, 10 min) and the supernatant was discarded. The pallet containing plates were frozen overnight at −20°C. The pallets were thawed, and lysed with 300 µL Lysis buffer (TAB, 1 mg/mL hen egg white lysozyme (HEWL), 200 µM PLP, 125 U/mL Benzonase) for 30 minutes at 37°C. The plates were centrifuged for 30 minutes (4800 rpm). 150 µL supernatant was mixed with 150 µL TAB containing L-serine (10 mM final), indole (5 mM final), and PLP (50 µM final) in 2-mL 96-deep well plates. The reaction mixture was incubated for 16 hours, 37°C, gently shaking. The next day, the reaction mixture was allowed to cool down, and 30 µL was mixed with 30 µL 2x aptamer sensor stock solution in low-binding microtiter plates (Corning, 2881). The fluorescent values were determined as described under the section UV-Vis spectroscopy, taken within one hour of incubation after which the aptamer sensor was thought to be degraded by endogenous nucleases from *E. coli* and the signal decreased.

### Chip fabrication

The microfluidic devices used for double emuslion generations (Supplementary Fig. 1) were fabricated following standard photolithography and soft lithography procedures using high-resolution acetate masks and SU-8 photoresist patterning.^49,50^

#### Photolitography

The microfluidic chips was designed using AutoCAD (Autodesk) and printed out on a high-resolution film photomask (Micro Lithography Services). The mask designs are published on dropbase (https://openwetware.org/wiki/Dropbase:_Double-emulsion-02). The master moulds of microfluidic devices were fabricated following standard hard lithography protocols. First, 15-µm-high microfluidic structures were patterned on 3” silicon wafers (Microchemicals) using high-resolution film masks and SU-8 2015 photoresist (Kayaku Advanced Materials) according to the guidelines of the manufacturer (SU-8 2000, Micro Chem). A MJB4 mask aligner (SÜSS MicroTec) was used to UV expose all the SU-8 spin-coated wafers. The thickness of the structures (corresponding to the depth of channels in the final microfluidic devices) was confirmed by measuring with a Dektat stylus profilometer (Bruker).

#### Soft lithography

The poly(dimethyl)siloxane (PDMS) mold was prepared by pouring a mixture of PDMS and curing agent in the ratio of 10:1 (w/w) over the Si wafer. The mold was then cured at 65 degrees overnight. The PDMS mold was cut out from the Si wafer to release the microfluidic chip. Next, the inlet and outlet holes were punched using a biopsy punch (1 mM). The chip was washed with dishwashing liquid and water, followed by ethanol (96%). The chip was dried using compressed air, and cleaned additionally by the application and removal of scotch tape. The glass surface on which the chip was to be bonded was cleaned with scotch tape. The microfluidic chip and glass surface were both placed in a Femto Diener plasma surface treater. A vacuum was created before flushing the chamber with oxygen. Next, the chips were treated with plasma. The chamber was quickly ventilated to retrieve the microfluidic chip and glass surface, and bonded by rolling the chip on the glass surface. The chip was baked at 65°C for 10 minutes.

#### Hydrophilic and hydrophobic treatment of the microfluidic chip

To apply the hydrophobic coating to the channels in the chip which creates the single emulsion (chip design https://openwetware.org/wiki/Dropbase:_Double-emulsion-02), the freshly baked microfluidic chips were flushed with 1% (v/v) trichloro (1H, 1H, 2H, 2H)perfluoroacetylsilane in HFE-7500 (3M-Novec). The chips were heated for 10 minutes at 75°C before storage at room temperature, sealing of the inlets and outlets with scotch tape. To apply the hydrophobic coating to the channels in the chip which creates the double emulsion (chip design https://openwetware.org/wiki/Dropbase:_Double-emulsion-02), the freshly backed microfluidic chips were first flushed with 0.2wt% poly(diallyldemethylammonium chloride) or pDADMAC in 0.5M NaCl. After 10 minutes, the channels were flushed with 0.1M NaCl, before flushing the chip immediately with 0.2 wt% poly(styrene sulfonate) or PSS in 0.5M NaCl. After 10 minutes, the chip was flushed three times with ddH2O, and kept submerged in ddH2O at 4°C until use.

### Microfluidics

#### Preparation of 2x cell solution

Two days prior to the w/o formation, the plasmid library or purified plasmid was transformed to *E. coli* C43 DE3 ΔTnaA and plated on LBkan-Agar plates. The next days, the cells were scraped with LBkan and diluted to OD600 of 0.8 in 20 mL LBkan in small Erlenmeyer flasks. The cells were induced with 1 mM IPTG final, and shaken overnight at 20°C, 250 rpm. The next day, the OD600 was measured. The cells were diluted to an OD600 of 0.84. This assumes an OD 1 to contain 2×10^8^ cells, and after 2x dilution with the substrate, sensor and lysis solution in a droplet volume of 6 pL, will result in a final occupancy of λ=0.5. The cells were pelleted (2000 g, 3 minutes) and washed 3x times with TAB.

#### Preparation of 2x substrate, sensor and lysis solution

A 4x aptamer stock solution (2 µM aptamer, 5 µM CS-10) was prepared as described above. A 4x substrate stock solution (40 mM _L_-Ser, 20 mM indole, 0.2 mM PLP) was prepared in a 2 mL HPLC glass vial, and incubated for 2 minutes at 65°C until all indole was dissolved after it was cooled down to room temperature. The 4x substrate stock solution was filtered through a 0.22 µM filter and mixed 1:1 with the 4x aptamer stock solution to the final 2x aptamer:substrate stock solution of 800 µL. Finally, 16 µL polymyxin (200 µM in 2x solution from 10 mM solution, 100 µM final in droplets) and 3.2 µL r-lysozyme (4 µL/mL in 2x solution, 2 µL final in droplets) were added.

#### Preparation of oil solution

HFE-7500 was filtered through a 0.22 µM filter (Millex) and mixed with RAN fluorosurfactant (RAN Biotechnologies) to a final concentration of 1% RAN. The solution was again filtered through a 0.22 µM filter (Millex).

#### Preparation and flushing of collection chambers

The collection chamber was a 0.5 mL Eppendorf tube, which was glued upside down on a glass plate. 1 mm in diameter holes were punched with a biopsy punch both at the top and on the side of the tube, nearer to the bottom. Polyethylene Portex tubing (0.38×1.09mm inner/outer diameter, SLS) or BOLA Tubing, PTFE (0.5×1.0mm inner/outer diameter) were attached with cyanoacrylate glue (PR1500, Scotch-Weld). The collection chambers were flushed with 0.22 µm filtered HFE-7500 (Novec 3M) prior to use, and filled with 1% RAN in HFE-7500. Droplets enter the collection chamber via the top, and the emulsion would float on top of the oil solution. Excess oil was discarded through the side channel at the bottom. For double emulsion formation, oil was pushed from the bottom channel, which pushed out the emulsion through the top channel to the microfluidic chip, where the droplets were encapsulated again.

#### Microfluidic rig, tubing and syringe setup

The microfluidic rig was a setup with neMESYS syringe pumps (Cetoni), and a high-speed camera (Phantom Miro eX2), which was mounted ona inverted light microscope (Brunel Microscopes Ltd.). Glass syringes (Hamilton), either 250 µL or 500 µL were used to contain the cell and substrate solutions. Glass syringe (SGE) of 2500 µL was used for the oil-phase. The tubing consisted either of polyethylene Portex tubing (0.38×1.09mm inner/outer diameter, SLS) or BOLA Tubing, PTFE (0.5×1.0mm inner/outer diameter). For the polyethylene tubing, Hamilton glass syringes were fitted with 26 gauge needles (0.464 mm outer diameter). For the PTFE tubing, Hamilton glass syringes were fitted with 22 gauge needles (0.718 mm outer diameter). For both sorts of tubing, the SGE syringes were fitted with a gauge 25 (0.515 mm outer diameter) needle.

#### w/o formation

The oil solution, 2x substrate, sensor and lysis solution, and the 2x cell solution were pumped with flow rates of 800/30/30 µL/h, respectively. This resulted in droplet sizes of ∼6 pL. Droplets were collected in the collection chamber as described above, with the tubing connected to the top of the collection chamber connected to the exit hole on the chip. Droplets were heat treated at 55°C for 1 hour to kill endogenous nuclease activity of *E. coli* upon cell lysis (*pf*TrpB is a thermophile and stable at 75°C). Next, the temperature was lowered to 37°C, and the droplets were incubated for a maximum of 72 hours.

#### w/o/w formation

TAB (1.5% Tween-20) and the w/o droplets were pumped with flow rates of 150/50 µL/h. A 250 µl glass syringe (Hamilton) containing 1% RAN (in HFE-7500) was fixed to the lower tubing of the collection chamber, and pushed out the droplets from the top of the collection chamber to the inner inlet. A small piece of tubing connected to the outlet channel collected the double emulsion droplets, and was placed in the middle of a 1.5 mL low binding tube (Eppendorf), which contained 1 mL TAB (1.5% tween), with the double emulsions settling at the bottom. The double emulsions were kept at 4°C until they were analyzed or sorted by flow cytometry/FACS, respectively.

### Flow cytometric analysis and FACS of double emulsions

Double emulsions were resuspended with a 200 µL pipet prior to measuring. Flow cytometric analysis was carried out on a CytoFLEX S machine for double emulsions stored in TAB (1.5% tween). Aptamer-Cy5 fluorescence was quantified using 640 nm excitation, with 660/10 nm bandpass filter. Flow cytometric sorting of double emulsions was performed on an ARIA III or ARIA II, with sorting into different low-binding tubes (Eppendorf) containing 100 µL nuclease-free water according to aptamer fluorescent intensity. Prior to sorting, the double emulsions were often diluted ∼5x in TAB (1.5% tween). Cy5 fluorescence was quantified using 633 nm excitation, with 660/20 nm bandpass filter. Importantly, the nozzle size to smoothly accommodate the 22.5 µm double emulsion droplets was 130 µM.

### Plasmid recovery

Immediately after sorting, 200 µL 1H,1H,2H,2H-Perfluorooctanol (PFO) (Alfa Aesar) was added to the ∼150 µL double emulsions in nuclease-free water, vortexed, and centrifuged quickly for 10 seconds. The top layer was extracted, and added to a DNA-low binding tube (Eppendorf). To the tube, 4 µL of UltraPure Salmon Sperm DNA solution (Thermo Fisher) diluted 100x in nuclease-free water (final 2500x dilution) was added. The leftover PFO with small amounts of aqueous phase on top was extracted once with a 100 µL solution of UltraPure Salmon Sperm DNA solution (Thermo Fisher), diluted 2500x in nuclease-free water. To the 200 µL recovered DNA, 1000 µL DNA binding buffer (Zymo) was added, and purified over silica columns (Zymoclean Gel DNA Recovery, Zymo Research), eluting in the minimal amounts of nuclease-free water. The resulting purified plasmids were transformed to *E. cloni* 10F ELITE Electrocompetent cells (Lucigen), and plated on two 140 mm Petri dishes containing LBkan-agar. The next day, the colonies were scraped, and the plasmid was isolated using Genejet plasmid miniprep kit (Qiagen). This purified plasmid stock was used for transformation to BL21 (DE3) competent *E. coli* (NEB, 2527) for rescreening in plates.

---

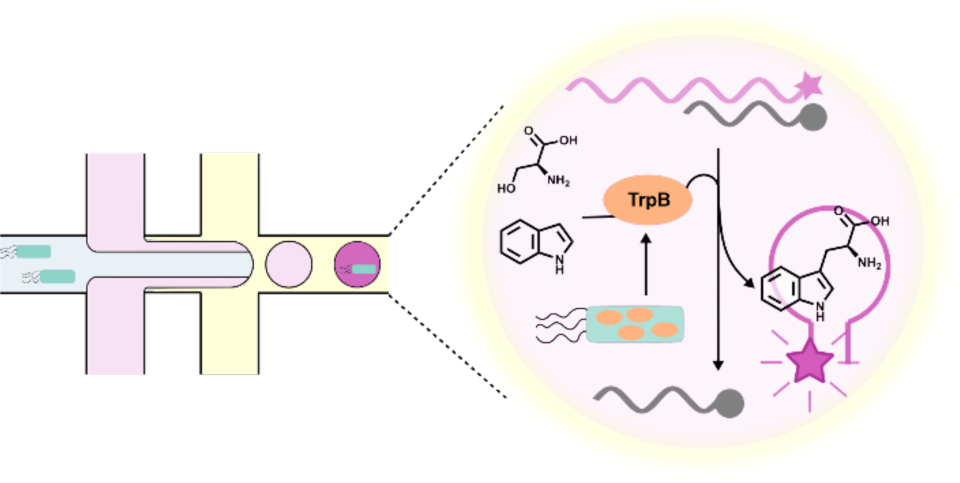
FOR TABLE OF CONTENTS ONLY

